# A High-Throughput Multiwell-Plate based Approach for the combined Expression, Export and Assay of Recombinant Proteins

**DOI:** 10.1101/2025.07.29.667353

**Authors:** Karen Baker, Daniel P. Mulvihill

## Abstract

High-throughput screening (HTS) of proteins has a wide range of applications across the biology, biotechnology, and medicine disciplines. These include yield optimisation, drug or biomarker discovery, and protein engineering, among others. Factors that need to be considered in designing high throughput protein expression and screening methods, be that for expression, activity, stability, or binding as says, include the required yield, reproducibility, solubility, stability, purity and activity of the protein. Thus, larger culture volumes and time-consuming manual protein extraction and purification steps are normally required to produce sufficient quantity of protein of appropriate purity. This limits the type of assay, and number of protein variants that can be simultaneously tested in an experiment. Here we describe a HTS protocol that allows the overnight expression, export and assay of recombinant proteins from *E. coli* cells in the same multi-well plate tube. The protocol uses a recently described Vesicle Nucleating peptide (VNp) technology that promotes high yield vesicular export of functional proteins from *E. coli* into the culture media. The resulting protein is of sufficient purity and yield that in can be used directly in plate-based enzymatic assays without additional purification. This simple single plate protocol allows itself to a wide range of high-throughput research and development screening applications, ranging from streamlining protein production and identification of activity enhancing mutations, to ligand screening for basic research, biotechnological and drug discovery applications.

## Introduction

High-throughput screening (HTS) is a tool used routinely in drug discovery and discovery research to enhance protein production and rapidly evaluate millions of compounds, molecules or proteins for activity against a biological target. It’s efficiency and scalability make it an attractive method for the rapid generation of large datasets, library screening, and for efficient optimisation of molecular design and expression of functional proteins. High-throughput protein expression and purification streamlines the rapid production and isolation of large numbers of proteins, thus reducing time and cost, accelerating both discovery research and drug discovery. Through application of simple, automated, scalable workflows, the approach enables the parallel processing and testing of multiple protein constructs, conditions, and strains, to optimise yield, solubility, functionality and purity. It accelerates the identification of suitable targets for biochemical assays, structural analyses, and therapeutic development. This approach is particularly attractive in screening protein variants, producing enzymes with enzymatic activities optimised for industrial applications. The ability to quickly produce high-quality proteins reduces time, cost and bottlenecks in research pipelines, while at the same time enhancing reproducibility compared to traditional methods. However current methodologies require extraction and purification of protein from cells requires that often require manual input to the process, and thus roadblocks to the HTS process.

We recently described the development of a Vesicle Nucleating peptide (VNp) technology ^1^ that provides an attractive alternative for the efficient production of recombinant proteins in the bacteria, *Escherichia coli*. The VNp tag facilitates the export of recombinant proteins into extracellular membrane-bound vesicles, creating a microenvironment that enhances the solubility and stability of challenging proteins, including those that are typically insoluble, contain disulfide bonds, or toxic to the bacteria. The VNp tagging approach not only increases protein yield but exports the protein into vesicles in a part-purified form that allows long-term storage of active proteins. It is thus a useful expression and downstream purification methodology, applicable to biotechnology, medical and fundamental discovery research applications alike.

Here we describe a rapid high throughput protocol using the VNp technology to express and isolate proteins from *E. coli* cells grown in multiwell format plates. The protocol describes rapid in-plate cold shock plasmid transformation, followed by in-plate protein induction and isolation, as well as an example subsequent in plate enzymatic assay. The protocol requires minimal sample transfer between plates and can be easily adapted and applied to a plethora of high-throughput research, development and library screening applications to streamline protein production for basic research as well as biomedical and industrial applications.

### Development of the protocol

*This protocol is a high throughput multiwell plate approach based upon the vesicle packaged recombinant protein technology, described in Eastwood et al, 2023 and Baker et al, 2025*.

While high export yields are obtained for many fusions using standard induction conditions ^1,2^, during the development of the Vesicle Nucleating Peptide fusion technology we found it necessary to optimise conditions to maximise the expression and export of the more challenging VNp-fusions tested. Several proteins required use of specific VNp fusion sequence variants, as well as different media, culture volume, induction levels, and promoters to obtain optimal yields.

To improve the throughput and enhance efficiency of expression and export of the VNp-fusion protein, we developed a multiwell-plate based protocol to allow a range of conditions to be tested simultaneously (e.g. volume, cell type, transcription induction levels, media, VNp variant, etc), to rapidly optimise growth, expression and export of the VNp-fusion. This screening method has been used successfully for construct design and optimisation of conditions to express, produce and package a range of functional proteins (e.g. IgG fusions, EPO, hGH and monoclonal antibodies) ^1,3^. The simplicity of the VNp expression and vesicular export system allows itself to rapid fusion protein isolation, with only a single centrifugation (or filtration) step required to isolate the fusion-protein filled vesicles from the *E. coli* cells. As the subsequent VNp-fusion protein within the vesicle fraction is relatively pure (typically > 85 % of total vesicular protein), and of high yield (typically ranges from 0.2 to 3 g/l from shaking cultures), it is well suited for integration into many HTS assay needs. This multiwell plate method was extended to facilitate in-plate enzymatic assay of the exported proteins, to identify optimal export and yield of functional protein, and implementation for an inhibitor screen. The uricase assay example included here can be easily adapted for use with other enzymatic reactions. If higher purity is required for the assays, in-plate affinity purification steps (also described here) can be easily integrated into the workflow.

During the initial development of the VNp technology it was found that surface area to volume ratio of the culture (cell oxygenation) is an important factor in the efficiency of vesicle formation and release ^2^. Critically, this ratio is more than sufficient for working volumes used in 24, 96, and 384 well format plate formats, that are routinely used in HTS methods. The protocols described here are optimised for 96 well format plates, which we use in this lab, as this allows sufficient working volumes and protein yields for subsequent analyses and assays we routinely undertake. The protocol can be used effectively with other well formats with appropriate culture volumes, as listed below. Users should select the appropriate well format according to the number of protein samples, quantity of protein(s) and volume necessary for their downstream needs.

### Applications of the method

While this HTS screen was developed for the rapid optimisation of protein export conditions and subsequent ligand binding / inhibitor screens, it can be applied to a wide range of applications including yield optimisation, expression libraries, drug or biomarker discovery, and protein engineering, among others. A simple application for this protocol, is to screen though a range of conditions to optimise expression and export of a protein of interest (Figure 1), by simultaneously culturing equivalent induction cultures in conditions. Only soluble, folded protein is exported into the vesicles (inclusions, when infrequently present, remain within the cytosol), the yields of folded protein can be determined using spectroscopy techniques. As we show here, the use of fluorescent protein fusions simplifies the analysis of protein expression and export (Figure 1). However, expression and export of non-fluorescent VNp-fusions with the Protein-of-Interest (POI) can be followed by SDS-PAGE analysis of the cell cleared media (Step 17) (Figure 2). Once conditions are optimised, expression and export for each VNp-fusion is extremely reproducible (Figure 2b), making it perfect for use in screening applications that require equivalent amounts of protein (or proteins). However, if purified protein is required for downstream analysis, the VNp-fusions can be rapidly purified using an in-plate affinity-purification protocol (Step 20) (Figure 2c).

**Figure 1:**
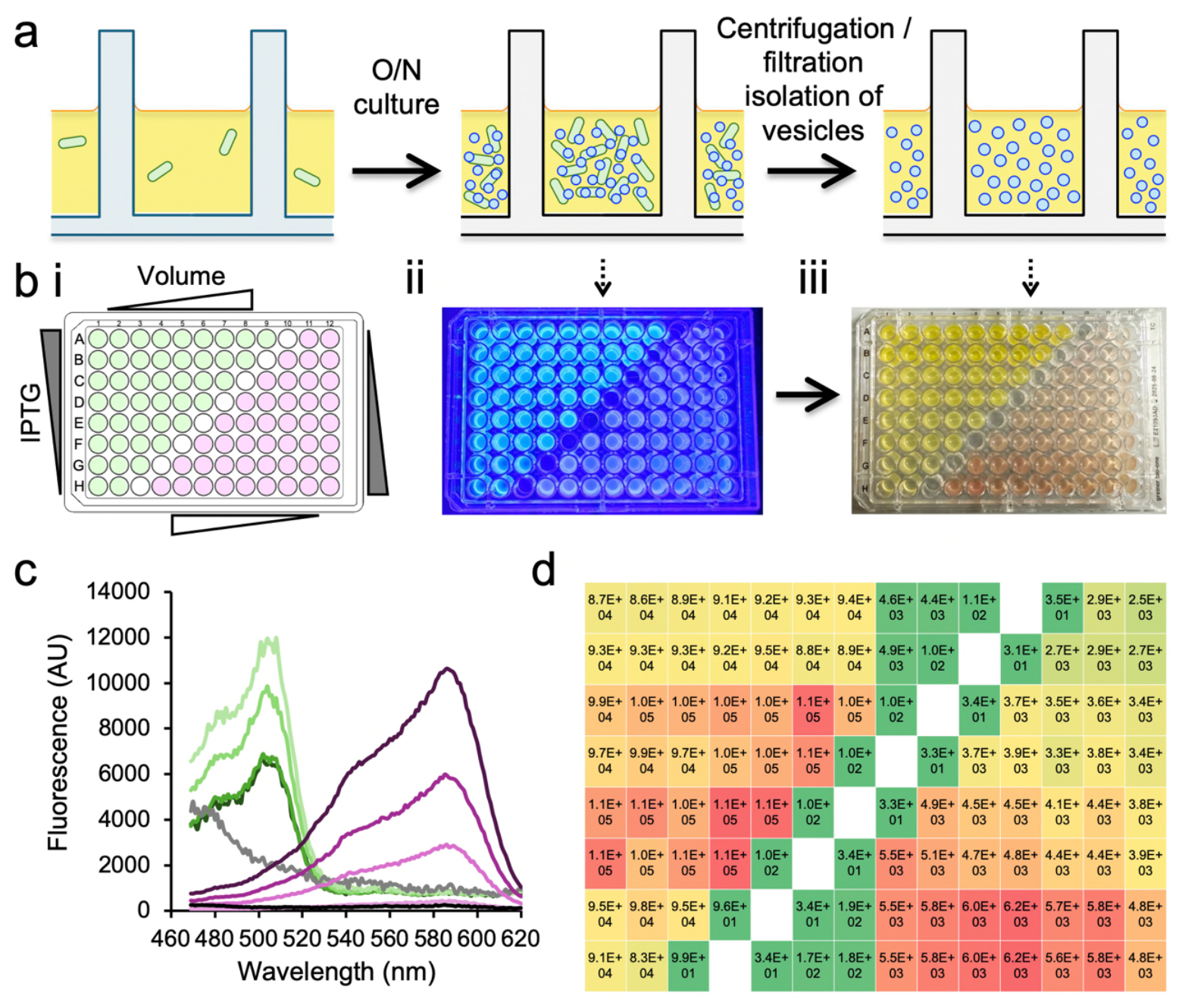
Multi-well plate format recombinant protein expression and vesicular export screen. (a) Schematic of multiwell plate format workflow for recombinant expression and isolation of fusion protein filled extracellular vesicles. (b) 96 well-plate expression and vesicle export optimisation screen of VNp-mNeongreen (green) and VNp-mCherry2 (pink). (i) Layout of VNp-mNeongreen (green) and VNp-mCherry2 (pink) expression construct containing *E. coli*, and test variables on the 96 well plate. (ii) Blue light illuminated in plate cultures (organised as set out in (i) after overnight induction. (iii) FP-filled-vesicle containing media from ii after cell removal by plate-well filtration. (c) Fluorescence scans of media from (Biii). Pink lines highlight VNp-mCherry2 abundance increases in proportion to culture volume (50 (light pink), 75, 100, 150 (dark magenta) µl). Green lines highlight variation in VNp-mNeongreen abundance in media from overnight cultures induced with 25 (light green), 50, 75, & 100 (dark green) µg / ml IPTG. (d) Heat map of mNeongreen (left) and mCherry2 (right) fluorescence in media from overnight 96 plate format cultures, as described in B.

**Figure 2:**
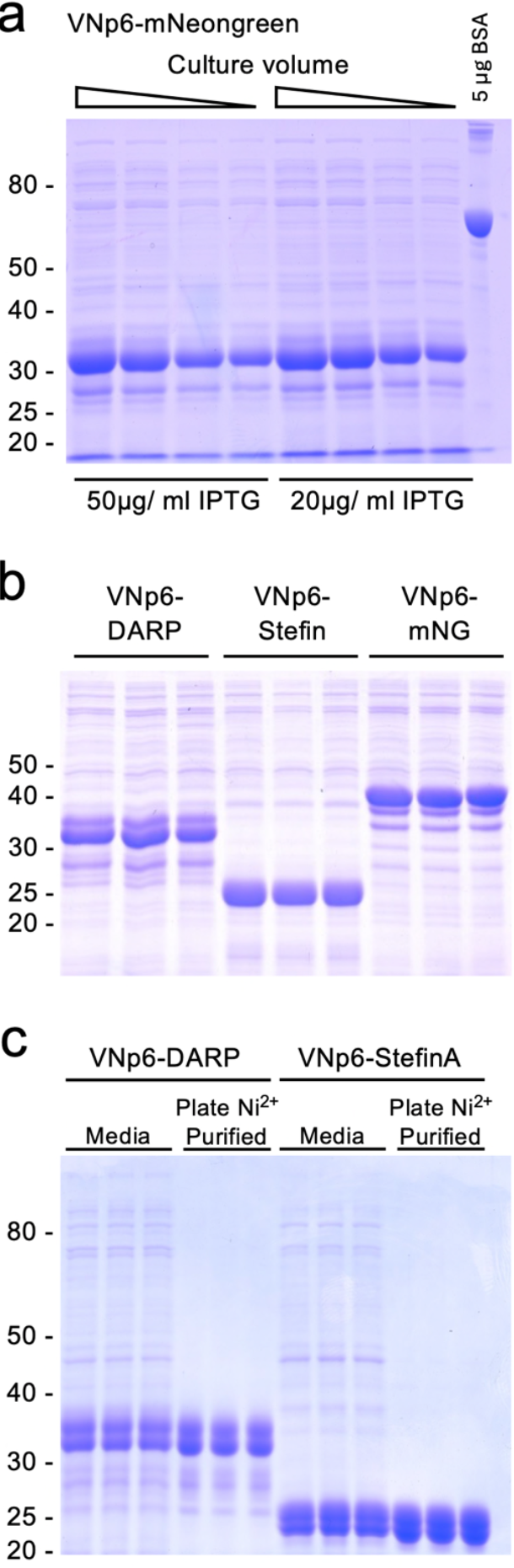
Examples of 96-well plate format vesicular protein export and purification screen samples. Commassie stained SDS-PAGE analysis of 10 µl media samples from different overnight 96-well plate cultures of *E*.*coli* expressing (a) VNp6-mNeongreen (samples from Figure 1), (b) VNp6-DARPin^off7^, VNp6-Stefin-A, or VNp6-mNeongreen fusions, highlighting yield enhancement (a) and reproducibility (b). (c) SDS-PAGE analysis of samples of media from 3 different overnight 96-well plate cultures of *E*.*coli* expressing either VNp6-DARPin^off7^ or VNp6-Stefin-A, as well as these carboxyl His6 tagged fusion-proteins purified from the same media samples using the described in plate Ni^2+^-NTA isolation (Step 20).

One such obvious HTS application is to directly follow enzymatic activity across samples within the plate. This is made possible due to the reproducibility, purity and yields (i.e. 40 to 600 µg of the VNp-fusion protein typically obtained from a 100 µl 96 plate well culture) of the vesicle exported protein (Figure 2) ^1^. As an illustration, we have developed a multiwell format *in vitro* assay that allows researchers to measure the activity of in-plate expressed and exported VNp-uricase protein (Figure 3), by following changes in 293 nm absorbance to monitor enzyme dependent breakdown of uric acid. The measured activities are extremely reproducible between individual culture wells, which is critical for high-throughput protein engineering screens. Thus by combining with a mutagenesis screen or mutant clone library, this protocol can be used to identify residues critical for protein function, and allows researchers to rapidly engineer proteins with enhanced activity, resilience and/or lifetimes for use in industrial and biotechnology applications. The yields allowed by this methodology also allow rapid biophysical and activity comparisons to be made of HTS protein variants of interest.

**Figure 3:**
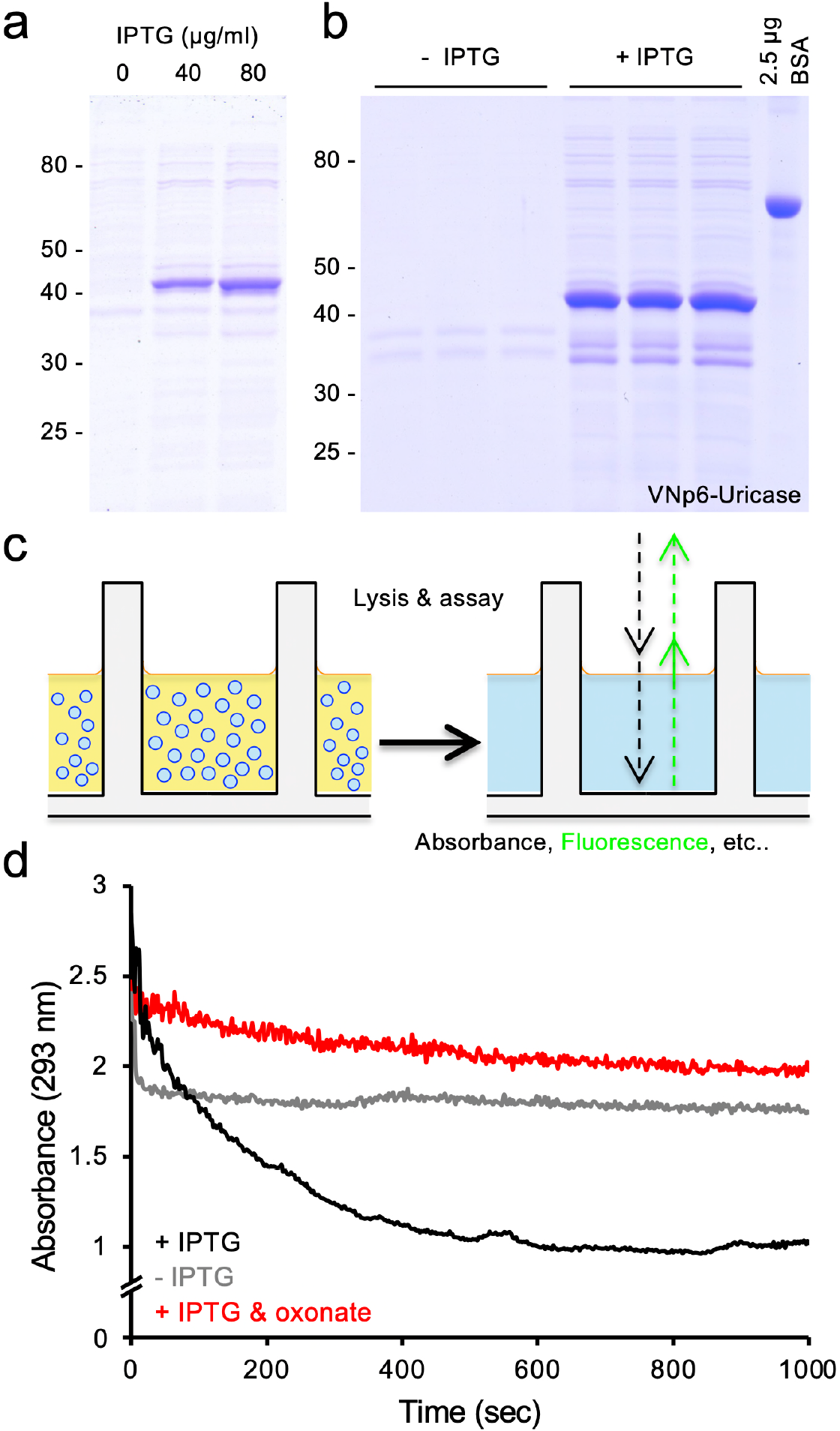
96-well plate format screen of vesicular export, activity and inhibition of active uricase. Commassie stained SDS-PAGE analysis of filtered 10 µl media samples from overnight 96-well plate cultures of *E*.*coli* containing a plasmid, expressing VNp6-uricase fusion (a) optimising IPTG concentrations, or (b) of uninduced (-IPTG) and induced (+IPTG) samples for subsequent in plate uricase activity screen. (c) Schematic of vesicle cleavage and plate-based assay. (d) 96-well plate uricase assays (curves averaged from 4 experimental repeats) in wells with media samples lacking uricase release (from b -IPTG sample) (grey line); with uricase released into the media (from b +IPTG sample) (black line); or samples with uricase released into the media with addition of 20 µM uricase inhibitor, oxonate (red line).

The high reproducibility of protein yields obtained using this protocol allows its simple integration into ligand binding screen assays. Ligands, which can vary from small molecules to peptides, can be identified that either bind to the protein of interest or inhibit / enhance its activity, as demonstrated by inclusion of the uricase inhibitor, oxonate, in the in-plate uricase assay Figure 3. The protocol can thus be used to efficiently screen small molecule drug libraries to identify protein effectors, and inversely, can also be used to express protein-variants, protein domain ^4^ or protein libraries ^5-8^ to screen for specificity and efficacy of an effector ligand. ^9^

The protocol is likely to be attractive for large scale drug screening applications, as the ability to be able to undertake high throughput in-well ligand binding and activity assays allows researchers to rapidly identify key functional and structural residues with a protein and thus gain mechanistic insights into its regulation and function. It provides a quick and cost-effective method to produce small quantities of a bank/library of proteins. By using the optional in-plate purification step (Step 20), allows the potential for the protocol’s application in high throughput crystallography screens. Finally, the ability to transform a wide range of strains using the cold shock steps described here, does away with the need to make banks of competent cells for each individual strain to be tested and allows the VNp-fusion expression and isolation protocol to be used to screen expression and activity of proteins across libraries of strains and genetic backgrounds.

### Advantages of the High-throughput Vesical Nucleating protocol over current methods

While a wide range of *E. coli* protein expression HTS have been described to date, ^10-13^ to our knowledge, this VNp based protocol is unique in that it represents the only HTS system allowing expression and vesicular export of a wide range of functional recombinant proteins from *E. coli* in a single step. ^14^ The protocol described here is unique as it allows expression, export and assay of a fusion protein in the same multiwell format plate. Normally, this protocol would require separate larger volume steps for (i) cell culture, protein expression and cell isolation; (ii) cell disruption; (iii) protein extraction; (iii) lysate clarification; (iv) protein solubilisation; and (v) purification in order to isolate sufficient protein of sufficient purity for either enzymatic activity or binding assays. ^15-17^ The VNp system has the additional benefits of not only allowing expression of more challenging proteins, but also expressing proteins at higher yields than standard methods will normally allow ^1^. This VNp based protocol allows all of the above stages to be completed in one plate step, as the protein is exported into the media within vesicular packages, in sufficient quantity and purity for a wide range of assays. Thus, the significantly increased yield, combined with doing away with the need for cell disruption and subsequent protein purification and concentration steps, allows simple integration into a rapid multiwell format HTS assay approach, that would not otherwise be achievable using any cell type.

While several eukaryotic expression systems promote export of recombinant proteins (e.g. *Pichia pastoris* or mammalian cell culture), ^18-23^ these systems can be technically more challenging to culture, take longer to grow, require more time for optimisation, and have higher consumable costs. In contrast, with this system typical optimised yields of exported VNp-fusion range between 200 mg to 3 g of per litre of culture. With standard working volumes on multiwell plates, these yields translate to typical yields of 0.2 to 3 mg, (24 well) 40 to 600 µg (96 well), and 16 to 240 µg (384 well) of exported, > 80 % purified protein. To our knowledge there are currently no equivalent cell or cell free systems or protocols available that are as quick, cheap and easy to use or able to achieve equivalent reproducible yields and purities in these HTS scales as the protocol described here.

### Experimental Design

#### Overview of the VNp method

The Vesicle Nucleating peptide technology requires fusion of a short amino-terminal amphipathic alpha-helix onto the protein of interest. This peptide interacts with the inner *E. coli* membrane, and at a critical concentration induces outward curvature of the lipid bilayer to form a vesicle which is released into the culture media. This strategy allows the protein to be packaged and released from the cytosol, thus allowing further protein production by preventing localised cytosolic accumulation of proteins that would otherwise impact the health of the cell. Due to basic differences in size and mass from the cells, the released vesicles can be easily isolated and transferred to a fresh plate by centrifugation. Once isolated, the vesicle can either be stored for > 1 yr at 4 ºC in the sterile filtered media, or membranes can be lysed by the addition of anionic (or zwitterionic) detergents and used directly in the downstream assay. This protocol requires only a single sample transfer step between plates, during the vesicle isolation, however a single fresh plate transfer will be required if the affinity purification of the protein is required. While we routinely use multi-pipettes when undertaking this high throughput multiwell plate protocol, the method can be easily adapted for integration into robotic liquid handling systems. Whatever detailed strategy and assay you decide upon, the initial design of the VNp-fusion construct is the most important factor for optimisation of this protocol.

#### Construct design and *E. coli* strain selection

We have generated and tested a large number of Vesicle Nucleating peptides, and for most applications would recommend using either the VNp2, VNp6 or VNp15 sequences ^1^ as these routinely result in highest yields for most proteins tested to date. When fused to the amino-terminus of a protein, these short (38 or 20 residues) peptide tags are sufficient to promote association with the membrane and subsequent encapsulation into a vesicle. While it may be attractive to fuse the VNp to the carboxyl terminus of some proteins, it is less likely to result in vesicular export using this strategy, and even when it does protein yields are significantly lower.

Protein and assay specific requirements for the design of the final fusion (e.g. affinity purification, solubility and fluorescence tags, labelling residues, spacers, etc) should be considered. While there is a lot of flexibility in the choice of backbone for the expression construct plasmid, there are two important constraints. First, the selection marker. Using an antibiotic marker that affects the bacterial cell wall (e.g. ampicillin) dramatically reduces the viability of cells when expressing VNp-fusion. While we routinely use kanamycin, chloramphenicol and tetracycline selection, all other non-cell wall antibiotics markers, tested to date do not measurably affect VNp-fusion export. Second, expression of the VNp-fusion must be under the control of a regulatable promoter as expression levels need to be repressed during transformation and be regulatable during expression to optimise vesicular export. To date we have used rhamnose, arabinose, and IPTG (both T5 & T7) inducible promoters, which together allow a range of options for repression and induction levels, as well as a wide choice of *E. coli* expression strains. The VNp system works best in B and K12 *E. coli* strains, and routinely use BL21 DE3 star cells for most VNp-fusion protein expression applications. However we have successfully used other *E. coli* strains when necessitated by the protein and / or assay.

#### Culture and induction

This protocol can be used to assay one plasmid (e.g. to identify optimal conditions) or multiple plasmids or multiple strains. If the former, a single starter culture should be used to inoculate each well culture, in order to maximise consistency between cultures. If the latter, individual starter cultures should be set up in each well. Most standard *E. coli* media can be used for this protocol, and while TB is routinely used to maximise both cell growth and protein expression, glycerol concentration should be kept to a minimum ^1^, to maximise vesicle formation. If minimal media is required for downstream labelling during the assay (e.g. N^15^ labelling for NMR applications), cells should be initially cultured in rich media and transferred to minimal media at induction (Step 12).

Expression should be induced at late log phase, at a typical OD600 of 0.8 to 1.0. Maximal induction of VNp-fusions using standard promoters leads to rapid cell lysis, and therefore recombinant protein transcription levels should be moderated. For example, IPTG should be used at a reduced final concentration of 20 µg/ml for most VNp-fusions expressed from genes under the control of the T7 promoter. It is worth noting that optimal protein production and optimal export conditions do not always coincide with each other, and therefore a balance needs to be struck to obtain maximal export of recombinant protein for the subsequent assay. For example, while vesicle formation is most efficient when cells are cultured at 37 ºC (lipid membranes are more dynamic), some proteins require expression at lower temperatures in order to fold correctly. In these situations, we tend to recommend using the VNp6 tag, that is able to promote membrane reorganisation across a wider range of temperatures.

One important internal control that should be considered when using this protocol, is to include control wells containing equivalent cells expressing a non-VNp labelled variant of protein to confirm VNp-dependent export and assay specificity. However, cell growth should be monitored during optimisation of the protocol to highlight optimal growth conditions and identify cell death brought about by overexpression of the VNp-fusion.

#### Vesicle isolation, protein release, and purification or assay

Harvesting vesicles requires separation from the cell culture and transfer to a fresh multiwell plate. This can be done easily by either centrifugation or filtration centrifugation. Cells can be pelleted with a 5 min spin, and the vesicle containing media can be carefully transferred to a fresh multiwell plate, appropriate for the assay (e.g. black wells, UV transparent, etc). Alternatively, the cultures are transferred to an appropriate multiwell filter that retains cells but allows vesicles to pass through. Whichever method is used, the plate containing samples of protein filled vesicles can either be stored long term at 4ºC at this stage, or used directly. Vesicles are lysed in plate by adding non-ionic detergent (e.g. Triton, Tween, etc) to release protein into the media. The protein can be assayed directly or purified using a simple in plate affinity purification protocol. The protein can then be used directly in any fluorescence or absorbance-based assay as required. We include the description of an enzymatic assay here to illustrate the functionality of vesicle packaged proteins and demonstrate how this protocol produces sufficient protein on multiwell plates to measure rate changes in signal with a HTS format. It is worth noting that the vesicle lysis step is only required if the assay substrate is unable to pass across a lipid membranes.

#### Limitations and development opportunities

Since first describing the VNp technology, we and others have found that it can be applied to a wide range of proteins from diverse organisms. Typically, if you can express your protein using standard expression and culture methods, the VNp method can be used and results in increased yield of exported functional protein. In addition, due to the compartmentalisation into membrane bound vesicles, many proteins that would otherwise be insoluble, toxic, or remain unfolded, can be expressed in a functional form using the technology (e.g. hGH, EPO, DNAse, monoclonal antibodies). In some cases, a varying proportion of the protein filled vesicles are internalised into the bacterial cytosol, thus requiring disruption of the cell wall before being used in downstream assays. A method for the simple isolation of these cytosolic VNp-fusion protein-filled membrane vesicles is currently in development.

In our experience, monomeric proteins of less than 85 kDa mass are most likely to be exported into extracellular vesicles, but this is not a hard and fast rule. Similarly, we find a proportion of dimeric proteins (e.g. Etanercept, leucine zipper fusions, etc) are often seen to be exported into the media ^1^, so it is worthwhile testing expression and export of the desired VNp-fusion using a flask culture as described in detail elsewhere ^2^, and then undertaking an expression and export optimisation screen, as described here (Figure 1). It is worth considering fusing your protein of interest with different amino terminal VNp tags ^1^ in combinations with mNeongreen, MBP, Sumo, or other solubilisation tags and to determine optimal construct for your application. Then screen (as described here Figure 1) to determine optimal combination of media, temp, induction level, induction timing, and culture volume for production of the protein.

## Materials

### Biological materials

- *E. coli* cells – the VNp tags work with a range of strains (see Eastwood et al, 2023), but to date, protocol works best with K12 and B strains, with BL21 DE3 Star optimal for most proteins we have worked with.
- Bacterial plasmid(s) for expression of VNp-fusions. Inducible promoter with non-cell wall targeting antibiotic selection (e.g. use tetramycin, kanamycin, or chloramphenicol). A range of constructs generated in this lab are available from Addgene (www.addgene.org/Dan_Mulvihill).

### Reagents

- Flat bottomed multiwell plates (e.g. Greiner # 655180)
- *V bottomed multiwell plates (e*.*g. Brand # 781661 or Sarstedt # 82*.*1583001) – use double orbital shaking during culture steps*.
- Gas permeable plate sealing film (*e*.*g. Brand # 701365*)
- Black UV transmission multiwell plates (e.g. *Brand # 781615*)
- Multiwell filter plates (*e*.*g. Whatman Unifilter Microplate #7700-3308*) – *Optional step 16*.
- Dry ice
- Industrial Methylated Spirits (IMS)
- Triton X-100 or Tween 20.
- Lysogeny / Luria-Bertani Broth (LB) media.
- Terrific Broth (TB) media.

### Equipment

- Shaking UV & fluorescence plate reader (e.g. BMG Labtech Clariostar plus) and/or shaking plate incubator (e.g. BMG Thermostar)
- Centrifuge with plate adapters (*e*.*g. Eppendorf 5920R # 5948000060 & 5895125000*)
- Multichannel pipette. (*e*.*g. Brand Transferpette S-8 # 705908*)

### Software

- Software associated with plate reader for operation and analysis

## Procedure

### 96 well-plate Transformation – Day 1

This first stage allows bulk cold-shock transformation to introduce a library of different constructs into the *E. coli* cells in a multiwell plate format. This allows a range of constructs to be simultaneously tested (e.g. use for comparing different promoters, VNp tags, fusion protein mutants/variants, strains, or different combinations of each).

NOTE: Recommend use single tube if only single construct and strain being used (use standard heat-shock method, as described in any standard laboratory manual, such as ^24^.

1. Inoculate wells each containing 100 µl of LB media with *E. coli* cells at 37 ºC using a shaking plate incubator / plate reader set at 200 rpm agitation (use double orbital setting if available).
2. Once the majority of cultures have reached an OD600 of 0.6 - 0.8, harvest the cells by centrifugation for 5 min at 2,250 x RCF) and aspirate off the media.
3. Resuspend in 50 µl of ice-cold 100 mM CaCl2 10% glycerol. Leave on ice for 10 min.
4. Add 1 µl of plasmid DNA from a 0.01 to 0.1 µg/µl stock to each well
5. Mix by agitation then place on ice for 10 mins.
6. Place onto a bath of dry ice and ethanol for 90 seconds NOTE: Put layer of dry ice at bottom of ice bucket and pour over with industrial methylated spirits (so base of plate is resting in liquid).
7. Immediately place at 37 ºC to thaw (typically 5 min).
8. Add 100 µl of LB and incubate for 1 hr at 37 ºC to recover.
9. Take ≥ 5 µl volumes from each well and grow transformants overnight using on the follow 2 methods: Option A:
  i. Plate out onto a matrix grid on an LB agar plate supplemented with appropriate antibiotic, and place in 37 ºC overnight. NOTE: Ensure LB agar plate has dried sufficiently to efficiently absorb culture aliquots.
  ii. Once colonies are visible in the morning, store plate at 4 ºC until ready to use.
  Option B:
  - Add to fresh multiwell plate with wells containing LB supplemented with appropriate antibiotic and culture overnight at 37 ºC with shaking. ***Total timing to this point: 3*.*5 hours (depending on growth rate in step 1)***. 10. Once colonies are visible in the morning, store the plate at 4 ºC until ready to use. PAUSE POINT: Plate can be stored for up to one week at 4 ºC.

### Expression, export and isolation of vesicular packaged recombinant protein - Day 2-3

11 Either pick single colonies (if followed 9A) or aliquot 5 µl from overnight culture wells (if followed option 9B) and transfer into wells containing up to 200 µl TB + antibiotic NOTE (Optimise the volume for your construct and downstream screen) – see Fig. 1.
12 Grow on heated shaking plate incubator (or preferably shaking plate reader with double orbital shaking) at 37 ºC, 200 rpm until cultures reach a late log phase density (typically less than 2hrs). Optional Step 1: Once cells desired reach density, change culture temperature at this point, if required for optimised protein folding. Optional Step 2: If protein expression with minimal media is required (e.g. for labelling applications), swap media at this point by
  i. Centrifuge at 2,250 x RCF for 5 mins
  ii. Remove LB media and replace with equivalent volume of warmed minimal media.
13 Induce protein expression by addition of transcription inducer molecule. NOTE: Typically use up to 50 µg / ml IPTG with T5 and T7 promoters. ***Timing to this point: Varies typically between 2 to 3 hours***.
14 Seal plate surface with gas permeable film to minimise evaporation of media.
15 Culture overnight at 37 ºC with 200 rpm shaking.
16 After at least 12 hrs, separate cells from media using one of the following methods: Option A:
  i. Centrifugation at 2,250 x RCF for 5 mins at 4 ºC.
  ii. Transfer supernatants onto required assay plate. Option B:
  i Transfer onto 0.45 µm filter plate mounted onto required assay plate.
  ii Centrifuge at 2,250 x RCF for 2 mins at 4 ºC. Vesicles and media will pass through to the fresh plate.
17 Place plate containing media (and exported vesicles) at 4 ºC. PAUSE POINT: Recombinant protein and vesicles can remain stable in sterile media for at least a year. ***Timing from step 16 to this point: 10 min***.
18 Lyse vesicle membranes by addition of non-ionic detergent (e.g. 0.1 % Triton-X100 or 0.1 % Tween 20). NOTE: If >80% protein purity is required), use in plate affinity purification (Step 20).
19 Sample is now ready for assay.

### In plate purification

20 Add 10 µl of affinity resin (e.g. Ni^2+^-NTA) to each media containing well (V-bottomed plates are optimal for centrifugation.
21 Mix on shaker at 4 ºC for 30 mins.
22 Centrifuge at 500 x RCF at 4 ºC for 2 min.
23 Wash in 100 µl wash buffer (20 mM Tris 500 mM NaCl 25 mM imidazole pH 8.0). Repeat steps 22 and 23 twice more.
24 Isolate protein from resin with elution buffer (20 mM Tris 500 mM NaCl 300 mM imidazole pH 8.0).
25 Spin and transfer the buffer, containing purified protein, to a fresh plate for assay.

### Example enzymatic assay – Day 3

Vesicles used in this example were isolated, using the method described above, from BL21 DE3 star *E. coli* cells containing the pRSFDUET-VNp6-uricase-His6 expression plasmid ^1^ (Addgene plasmid #182401).

26. Add 50 µl of 2 mM uric acid (dissolved in 100 mM Tris (pH 8.5)) to UV transmission flat bottomed 96 well plate.

27. Using a plate reader, measure absorbance at 293 nm every second at 25 ºC for 1 min.

28. Add 50 µl of exported protein sample from step **18**.

NOTE: can add inhibitors / test inhibitors to samples at this point.

29. Immediately continue monitoring absorbance at 293 nm every second at 25 ºC until end of reaction in all wells.

***Timing from step 20 to completion of assay is typically 10 min***.

## Acknowledgements

The authors thank S. Cuce, B. Streather and T. Eastwood for testing and providing feedback on the protocol. This work was supported by the University of Kent and funding from the Biotechnology and Biological Sciences Research Council (BB/S005544/1 & BB/X007448/1). The Vesicle Nucleating peptide technology described here is described in the international patent filing *PCT/GB2022/053239*.

## Author contributions

KB performed the laboratory experiments; KB and DPM designed experiments; DPM sought funding and supervised the project. DPM wrote the main drafts of the manuscript and both authors contributed to editing.

